# Multiomics analyses reveal the central role of nucleolus and nucleoid machinery during heat stress acclimation in *Pinus radiata*

**DOI:** 10.1101/2022.07.08.499117

**Authors:** Mónica Escandón, Luis Valledor, Laura Lamelas, Jóse M Álvarez, María Jesús Cañal, Mónica Meijón

## Abstract

Climate warming is causing quick changes in mean annual temperature and more severe drought period. These are major contributors of forest dieback, which is becoming more frequent and widespread, particularly in warm and drought-prone regions. Despite being a hot topic in non-woody plant sciences, the information about how heatwaves impact in tree molecular biology is still scarce. In this work we investigated how the transcriptome of *Pinus radiata* changes during initial stress response and stress acclimation. To this end, and considering this species is non sequenced, we generated a deep dataset employing Illumina technology. This approach allowed us to reconstruct 77335 contigs which were annotated following gene ontology, and to define 12164 and 13590 transcripts as down- and upregulated, respectively, across the three sampled experimental points. Enrichment analysis allowed to distinguish 9 down-regulated pathways, the most of them related to the reduction of apoplast, and water transport. While 22 were upregulated, which followed two different trends those pathways that peaks at short-term (acute response) from those which accumulated long-term (acclimation response) being most of them related to heat shock response, redox machinery and RNA processing. Additionally, the combination of transcriptome data with other available omics layers, allowed an exceptional understanding of the mechanisms behind heat stress response, involving complex interrelated processes from molecular to physiological level. Nucleolus and nucleoid activities seem to be a central core in acclimating process, producing specific RNA isoforms and other essential elements for anterograde-retrograde stress signaling as NAC proteins, Helicase RVB, RZ1 RNA chaperone, or ribosomal RPS4. These mechanisms are connected by elements already known in heat stress-response (redox, heat shock proteins or ABA-related). But also, novel candidates, as photosynthetic pigments, shikimate, or proline centric proteases activities, have been identified effectively networking biochemical responses to its potential regulatory element. This work provides a first deep overview about what molecular mechanisms underlying heat stress response and acclimation in pines, supporting the development of new breeding strategies to face the challenges that the climate change will impose to forests.

## Introduction

According to numerous climate models and records, temperatures are rising, and heat waves events are increasingly frequent. The climate change framework coupled with the increasing wood demand (FAO, 2009) highlights the importance of the study of adaptation mechanisms of forest largely dependent on climate and is highly vulnerable to climate change (Wahid et al., 2007).

Like in other plant species, heat stress (HS) causes in Pines a detrimental and irreversible damage affecting development, growth, and in last term a population decline in natural forests or productivity losses in plantations (Ferretti et al., 2021a, 2021b; Gazol et al., 2020). A variety of responses to minimize damage and ensure protection of cellular homeostasis have been identified and significant progress has been achieved in the elucidation of the physiological and molecular mechanisms underlying (Escandón et al., 2017; Kotak et al., 2007; Li et al., 2021). However, despite some signaling mechanisms mediating heat stress acclimation have been described in *Pinus radiata* (Lamelas et al., 2022), there remain some gaps that need to be addressed, particularly those regarding the specific transcriptional regulation in response to heat stress and its integration with specific signaling mechanisms and the physiological response.

Plants and trees have developed excellent stress perception mechanisms and signal transduction. In response to heat stress, a multitude of processes are activated which enable the plants to cope with these stresses up to a certain extent. Some studies have suggested that plants developed complex regulatory networks at the biochemical, physiological, and transcriptional levels to cope with HS (Yan et al., 2020). The first response of plants to HS involves an increase in membrane fluidity of the plasmalemma. This leads to the production of secondary messengers and chemical signals such as calcium in the cytoplasm. Mitogen-activated protein kinases (MAPKs) and calcium-dependent protein kinases (CDPKs) are in turn immediately regulated by the Ca^2+^ influx. To maintain the water balance and osmotic adjustment in the cell, these signaling components are further transmitted to the nucleus, resulting in the induction of antioxidants, electron transport and solutes that function osmotically in the cytoplasm. If the stress persists, acclimation required deeper changes such as the accumulation of heat-tolerant protein isoform or the modification of membranes, is the result of a precise proteome remodeling (Escandón et al., 2017) finely tuned by specific signaling networks (Lamelas et al., 2022).

Recent studies in *A. thaliana* (Ohama et al., 2017) elucidated small portions of complex transcriptional networks during HS which revealed the complexity of the response. In *P. radiata*, previous works demonstrated that heat shock proteins (HSP), hormone related proteins and reactive oxygen species (ROS)-scavenging enzymes are the major functional proteins induced by heat stress (Escandón et al., 2016, 2017, 2018), however the molecular mechanisms leading to this remodeling are still unknown. Post-transcriptional regulatory mechanisms involving the processing of precursor mRNA, such alternative splicing, and other proteins related to RNA metabolism are proposed to play an important role in HS response (Lamelas et al., 2020, 2022; Ling et al., 2018), but there is no clue about how these regulatory mechanisms are integrated to coordinate a systemic response.

One of the most important open questions about HS response is how plants sense high-temperature and transduce this signal to transcriptional level. At least four putative sensors have recently been proposed to trigger the HS response. They include a plasma membrane channel that initiates an inward calcium flux, a histone sensor in the nucleus, and two unfolded protein sensors in the endoplasmic reticulum and the cytosol (Ohama et al., 2017). Each of these putative sensors is thought to activate a similar set of HS response genes leading to enhanced thermotolerance (Mittler et al., 2012). The transcriptional regulation of a thermotolerance program is, at the end, a convergent pathway in which different triggers can induce changes at chromatin and RNA processing levels. However, multiple, complex, and dynamically intertwined interactions among transcripts, proteins, and metabolites determine the final response of plants towards HS (Ohama et al., 2017). Multi-omic analyses can be considered a good approach for addressing complex biological interactions, as it has been demonstrated as a power tool to reveal different regulatory units across diverse omics layers (e.g., obtained from RNA, proteins, metabolites, etc.) (Escandón et al., 2017; Pascual et al., 2017; Valledor et al., 2021; Zhao et al., 2020). In last years, multi-omics efforts have taken center stage in basic research leading to the development of new insights into complex biological events and processes (Krassowski et al., 2020).

The aim of this work was to analyze the variations on the needle transcriptome during a HS acclimation in *P. radiata*, aiming to understand these changes based on different molecular interactions with other regulatory and physiological layers. The discovery of these mechanisms is always complex, and even more in unsequenced and poorly annotated species such as *P. radiata*. To increase the biological significance of this work we applied a multi-omics approach combining the newly generated transcriptome with available datasets for metabolome, proteome and physiology which were generated under the same experimental conditions (Escandón et al., 2016, 2017, 2018). This work provides new insights into the response to HS, pointing the central role of the posttranscriptional processing of RNA in nuclei and the different organelles, and how it is connected to cell, tissue, and physiological acclimation through different heat stress response mechanisms (red-ox-, HSPs-, ABA-mediated), suggesting an important function of shikimate pathway. Overall, this work provides new insights and propose new mechanisms of HS acclimation that increase our understanding of how plants adapt to challenging environments. In the current context of widespread tree dieback, defining the molecular basis of tree resilience capacity needs to be addressed to be able to develop tools for increasing the efficiency bred programs against climate change threats.

## Material and Methods

### Experimental design and RNA extraction

RNA was extracted from mature needles belonging to a time-courses heat-stress experiment (Escandón et al., 2017). In brief one-year-old *P. radiata* seedlings (plant size about 33 ± 4 cm) were kept under controlled conditions (Fitoclima 1200, Aralab) with a photoperiod of 16 h (400 μmol m^-2^ s^-1^) at 25 °C and 50 % relative humidity (RH), and 15 °C and 60 % RH during the night period. Before starting the experiment, a group of control plants (C) were collected. Heat exposure treatment began with a temperature ramp ranging from 15 to 40 °C over 5 hours and then hold at 40 °C for 6 hours and then ramped down to night values. The stress was applied for 5 consecutive days, and the plants were sampled after the end of 40°C period after first (T1), second (T2), third (T3) and last day (T5) (Figure S1). Three biological replicas (a pool of three plants each) were sampled at every timepoint. 100 mg of tissue (fresh weight) were ground to a fine powder in liquid N_2_ and RNA was extracted employing a modification of Valledor et al. (2014). The modification consisted of adding pellet solubilization buffer (PSB) directly to plant material omitting metabolite isolation steps.

RNA quantity of Control, T1 and T3 samples was checked using Qubit 3.0 Fluorometer (Thermo Fisher Scientific, USA) and integrity was assessed in a 2100 Bioanalyzer system (Agilent, USA) considering only samples with a RIN number above 6.9.

### RNA-Seq and RT-qPCR

#### cDNA library construction and sequencing

mRNA enrichment and cDNA library construction were conducted employing Illumina’s TruSeq Stranded mRNA Library Preparation Kit from total RNA and the sequencing of cDNA molecules was performed on Illumina Hiseq 2500 platform, using 100bp paired-end sequencing reads according to the manufacturer’s instructions (Illumina, San Diego, USA). This analysis was performed at StaBVida NGS facility (Caparica, Portugal) over Control, T1 and T3 samples (three biological replicates at each time). Every raw sequence data underwent adapters and low-quality bases trimming. The resulting reads were reanalyzed using SeqTrimNext software (Falgueras et al., 2010). Only the pairs in which both reads passed the quality test were further analyzed. The raw RNA-seq data have been deposited with links to Bioproject accession number PRJNA851373 in the NCBI Bioproject database (https://www.ncbi.nlm.nih.gov/bioproject/).

#### Transcript reconstruction from RNA-seq

All reads were trimmed employing TrimGalore (v0.6.5, Krueger, 2015) and ribosomal reads removed employing bowtie2 (v2.4.1, Langmead and Salzberg, 2012) and SILVAdb (release 102, Quast et al. 2013). Remaining reads were processed with trimmomatic (v0.32, Bolger et al. 2014), also discarding lectures shorter than 31 bp. Trinity (v2.10.0, Haas et al., 2013), denovoSOAP (v1.05, Xie et al. 2014), Trans-Abyss (v2.0.1, Robertson et al. 2010) and velvet/oases (v1.2.1.10/v0.2.9, Schulz et al. 2012) were employed in parallel for transcriptome reconstruction. Trinity was used employing default settings. For the other algorithms different kmer sizes were tested (26 to 64) and results were merged. Redundant sequences were removed employing cd-hit to each assembly (v4.8.1, Li and Godzik, 2006). The four generated transcriptomes were evaluated employing BUSCO (v4.1.2, Simão et al. 2015) and the best results were obtained after merging all of the obtained contigs. This combinatorial database was simplified removing redundant or shorter sequences by using evigene (Gilbert, 2019). A total of 77335 contigs were defined. These contigs were used as an in-house database for protein identification (see below).

#### Reads mapping and transcript quantification

Trimmed and filtered reads were mapped to the merged assembly using bowtie2 (v2.4.1, Langmead and Salzberg, 2012), and then independently quantified using Salmon (v1.2.1, Patro et al. 2017). Mapped reads represented the 97.97 %, 97,32 % and 97.64 % of the paired reads in control samples; 97.85 %, 96.94 % and 97.38 % in T1 samples and, 98.00 %, 97.24 % and 97.58 % in T3 reads.

#### Transcriptome annotation

Transcriptome sequences were locally annotated by using Sma3s v2 (Casimiro-Soriguer et al., 2017) and Uniref90 as reference database, and against Pfam database using dammit (v1.0, Scott, 2016) tool. Sequences were also classified according GeneOntology using GOMAP (V1.3.7, Wimalanathan and Lawrence-Dill, 2021), to MapMan by using Mercator4v2 (Schwacke et al., 2019). Out of the 77335 contigs, 48800 contigs were annotated, which averaged the 96.69 % of the sample’s quantification.

#### Targeted transcriptomics RT-qPCR

One μg of RNA was used to obtain cDNA using the RevertAid kit (Thermo Scientific, United States) and random hexamers as primers following the manufacturer’s instructions. CFX Connect Real Time PCR machine (Bio-Rad) was used to qPCR reactions with SsoAdvanced Universal SYBR Green Supermix (Bio-Rad, United States); three biological and two analytical replicates were performed for each treatment. ACTIN (ACT) and GLYCERALDEHYDE 3-212 PHOSPHATE DEHYDROGENASE (GAPDH) genes were used as endogenous genes as previous publications (Escandón et al., 2018; Lamelas et al., 2020). Normalized Relative Quantities (NRQ) and Standard Errors of RQ were determined according to Hellemans et al. (2007). Primers were designed using transcript sequences from RNA-Seq data. Detailed information about the primers used for qPCR experiments is available in Table S1.

### Protein identification and quantitation

Orbitrap Raw files already generated (Escandón et al. 2017; PRIDE identifier PXD032754) were revisited for employing the transcript database generated in this work for protein identification and quantification. Transcripts were six-frame translated and all peptides greater than 75 aa were kept. Proteome Discoverer 2.1 (Thermo Fisher Scientific, USA) was employed for re-analyzing files. Sequest HT and MS Amanda algorithms were employed with an FDR of 5%. The variable modifications were set as oxidation of methionine and protein N-terminal acetylation. Quantification was made with precursor ion area detection where protein peak areas were calculated on the basis of the top six unique peptides for the given protein.

### Statistical analyses

#### Data pre-process, univariate and integrative analyses

Transcriptomic data was filtered in order to keep only annotated contigs. In addition, available proteome and metabolome (Escandón et al., 2017; MTBLS4567 MetaboLights study, Haug et al. 2020), and hormonal and physiological datasets (Escandón et al., 2016) were employed in these analyses (Tables S2 and S3).

All statistical procedures were conducted with the R programming language running under the open-source computer software R v4.0.1 (R Development Core Team, 2020) employing RStudio v1.1 IDE (RStudio Team, 2016). All omics datasets were pre-processed and then analyzed with an integrated multivariate analysis employing pRocessomics v0.13 (https://github.com/Valledor/pRocessomics/) R package. All omics datasets were firstly pre-processed (missing values were imputed, abundance balanced, and z-score transformed). Univariate ANOVA test and q values were performed. PCA were performed over featured datasets in which non-variable variables, ANOVA p-values greater than 0.3, and the 15% of variables with higher coefficients of variation were filtered out. Multiblock sparse partial least squares-discrimant analysis (MB-sPLS-DA) (Singh et al., 2019), a data integration analysis for biomarker discovery using latent variable approaches for omics studies, was used to data integration. The optimal number of variables selected in each dataset was set in 60 transcripts, 50 proteins, 30 metabolites and 5 physiological parameters per component after tuning the function parameters employing the automated function available in mixOmics package (Rohart et al., 2017). Three biological replicates of each omics level were employed.

#### Pathway and enrichment analyses

Pathway analysis was conducted in parallel employing GO classification. Abundances were visualized employing heatmap representation. Enrichment analyses were performed employing g:Profiler with custom GMT annotation filse parsed from GOMAP following the recommendations provided by Reimand et al. (2019).

#### Network analyses and visualization

Networks obtained after MB-sPLS-DA analyses were analyzed in Cytoscape 3.9 (Cline et al., 2007) employing the plugin StringApp (Szklarczyk et al., 2017). STITCH network was constructed using *Arabidopsis thaliana* STRING database (only proteins with identity cut-off > 50%) and STITCH compounds included in the MB-sPLS-DA selected KeepX.

## Results and discussion

### Transcriptome profile of *Pinus radiata* needle response to heat stress

To investigate the gene expression dynamics in Pinus needles during the initial stages of heat stress response, first we assembled all reads to obtain 77335 contigs which were then employed to map reads corresponding to individual samples. From these sequences, we defined the best models for reducing redundancies, 48K dataset. On average 30.2M paired-ended reads were obtained per sample and a 97.55 % of the reads were mapped to contigs. These contigs were identified by homology employing Uniref90 database and Sma3s tool annotating a total of 45833 sequences. Mercator and GOMAP tools allowed respectively the functional classification of 36416 and 44789 contigs according to Mapman Classification and Gene Ontology (Figure S2).

For a global comparison of the transcriptomes of the 9 samples we initially performed a principal component analysis (PCA). To reduce noise and increase the meaning of the analyses, first we subtracted from the dataset non-variable and noisy variables. This featured dataset, counting 29602 transcripts (Table S1), demonstrated to be useful to distinguish the different treatments. The first two principal components (PCs) accounted for 68.2 % of the total variance (Figure 1A; Table S2). PC1, gathering 40% of the total variance, separated the different samples according to stress time. Transcripts exhibiting positive higher loadings to PC1, and thus explaining stress acclimation, included several RNA interacting and ribonuclear proteins and epigenetics-related proteins, characteristic of processes involved a transcriptome reprogramming, stress-signalers such as SNRK2 (Colina et al., 2020) and other kinases, cell cycle proteins, and solute transporters, which accumulation probably aims to retain water and to avoid a heat-driven drought/hyperosmotic cell stress. On the other hand, the proteins negatively correlated to PC1, and characteristic of control plants, pointed to normal plant metabolism and respiration, with no striking enrichment of transcripts belonging to any pathway. PC2 accounted for 28 % of the total variance and represented the initial thermal shock (grouping C and T3, and T1). Transcripts showing positive correlation to this component included some of the proteins correlated to PC1 (positively and negatively), explaining the homeostatic status of Control and T3 plants, as solute transporters or some heat shock proteins, however new signaling related transcripts (LRR-ROCO, F-BOX, WNK1, etc), fatty-acid and membrane metabolism, secondary metabolism of phenolic compounds or callose biosynthetic machinery appeared, probably indicating a subjacent drought stress. Proteins exhibiting higher negative loadings to PC2, correlating to T1 samples, suggested a “heat-shock” status with increased stress signaling mechanisms mediated by ROS-peroxisome (peroxin accumulation; Su et al. 2019), ethylene, nitric oxide, and other sensing proteins (ethylene response transcription factor and NO synthase (Hasanuzzaman et al., 2013), TIR-NBS-LRR (Kim et al., 2015), SnRKs proteins) and high number of heat shock proteins and several kinases. Transcripts also suggested, as in PC1, a proteome remodeling as ubiquitination mechanisms and RNA processing related proteins were also upregulated at T1.

**Figure 1:**
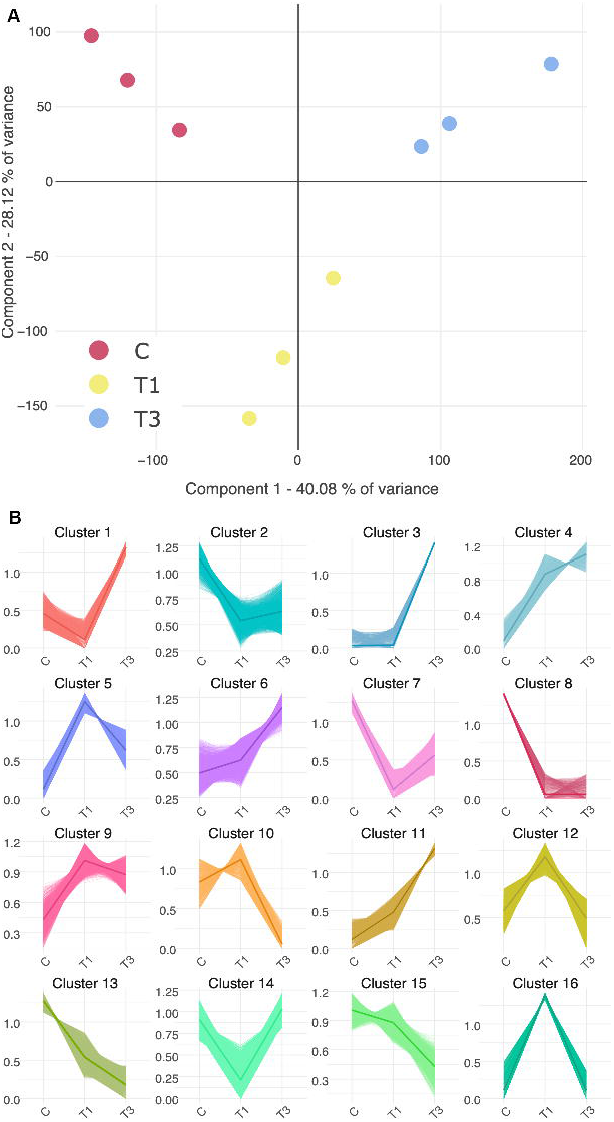
**A**. Sample classification after PCA analysis of transcriptome dataset. **B**. Sixteen distinct clusters of transcripts expression profiles in control (C) and after one (T1) and three (T3) days of heat-stress obtained after k-means analysis (Table S3). Thick lines represent the average expression of elements of each cluster.

Gene expression clusters were revealed after a k-means analysis (Figure 1C). Among clusters, the most recurrent trend was an overexpression T1 followed by a reduction in expression levels in T3 (clusters 5, 9, 10, 12 and 16). This type of clusters includes contigs related to heat stress (*HSP20, HSP70* and *HSP90*), but mostly owning to family HSP20 (17.5 kd *HEAT SHOCK FAMILY PROTEIN, MITOCHONDRIAL* and *CHLOROPLAST SMALL HEAT SHOCK PROTEINS*; Table S3) and genes involved in redox reactions, transcriptional regulators and ribosomal proteins (*TAU CLASS GLUTATHIONE S-TRANSFERASE*, mTERF1 or 60S ribosomal protein L36). Five clusters gathered those genes that drastically (2, 7 and 8) or gradually (13 and 15) downregulated, where genes included in lipid catabolic process (*GDSL ESTERASE/LIPASE*) or transport of water (Plasma *MEMBRANE INTRINSIC PROTEIN*) were overrepresented. Clusters 4, 6 and 11, grouped those genes which expression increased during the experiment, mainly related to in defense/stress response (*PUTATIVE TIR-NBS-LRR PROTEINS*), *RETROTRANSPOSON* genes (*RETROTRANSPOSON PROTEINS, PUTATIVE, TY3-GYPSY SUBCLASS*), and transcriptional regulators.

### Enrichment analyses distinguished short-term stress responsive- and acclimation-related pathways

The employ of Gene Ontology (GO) annotations allowed a comprehensive functional analysis of differentially expressed genes. Differentially expressed genes were mapped to GO and potential functional enrichments were revealed employing g:Profiler (Figure 2A). A total of 12164 and 13590 transcripts were considered down- and upregulated (±2-fold, p < 0.05, t-test of T1 and T3 times versus control) (Figure 2B). Among the 9 downregulated pathways, the reduction of apoplast, water transport, and cell wall related pathways may explain the observed increase of membrane damage and electrolytic leakage observed in this experimental system (Escandón et al., 2016). Interestingly, the downregulation of light harvesting and photosystem I related genes and heme binding proteins did not impact overall Fv/Fm performance, despite pigment biosynthetic enzymes such as porphobilinogen synthases (Table S4), or precursors such as heme (Table S5) are down accumulated. Lipid binding genes were also downregulated, being these results in the same line than those variables outlined by PCA and k-means analyses and supporting the sources of variation gathered by principal components.

**Figure 2:**
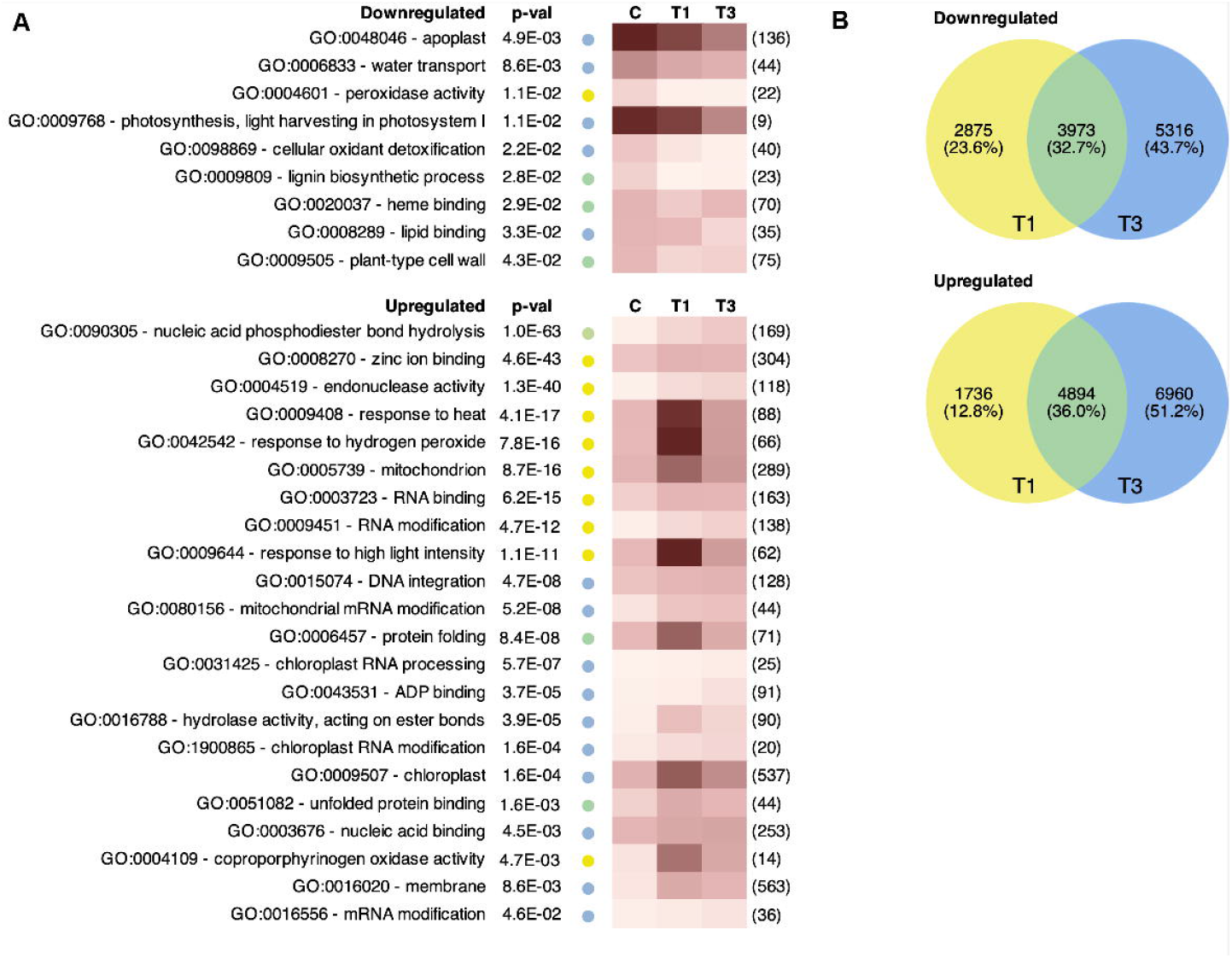
**A**. GeneOntology (GO) enrichment analysis of differentially expressed genes in response to heat stress at the two sampling times (T1, T3) with respect to controls (C). p-val column indicates the p-values obtained for each term after g:Profiler analyses. Colored dot indicates if term was enriched at T1 (yellow), T3 (blue) or both (green) sampling times. The color of the boxes represents the average expression rate of all differentially expressed genes (p < 0.05; >2-fold change) within each category at each sampling time. The number of differentially expressed genes within each category (TnQ value) is in brackets. **B**. Venn analysis of up and down regulated genes in T1 and T3 compared to controls.

On the other hand, 22 pathways were upregulated. Contrary to downregulated pathways, in which most of them followed a progressive decrease in their abundance, upregulated pathways followed two different behaviors, those pathways that peaks at T1 (acute short-term response) from those which accumulated long term, peaking at T3 (acclimation response). Initial responses seemed to be intended to reduce the heat impact and the initial protein denaturation and oxidative stress (response to heat, protein folding, high light, H_2_O_2_, chloroplast, and mitochondria) (Escandón et al., 2017). Longer term responses involved RNA processing at the different organelles and DNA related pathways, indicating the importance of the chromatin remodeling and the importance of producing new transcriptional isoforms with new activities or more stable protein forms for acclimating to heat stress (Lamelas et al., 2020). In Arabidopsis, new isoforms at chloroplast and mitochondria decrease the production of free radicals by adapting electron transfer chain complexes while keeping organelle function and energy production (Pascual et al., 2021). Pines seems then to behave in the same way, as at T3 photosynthetic rates (Fv/Fm) is recovered and auxin concentration, known modulators of electron transfer chain rearrangements (Berkowitz et al., 2016; Tivendale and Millar, 2022), increased, and light excess- and stress-related pathways are downregulated in comparison to T1. The integration of other omic levels with the transcript dataset, supported previously raised hypothesis and gave a broader functional perspective, benefiting from the synergies and molecular cross-validations that can only be reached when using multiple data layers, as is detailed below.

### The employ of an experiment-specific transcriptome database allowed a deeper proteome analysis which unveiled the role of nucleoids and RNA processing in stress acclimation

The available raw files of the proteomic analysis of this experimental system (Escandón et al., 2017) were reprocessed (identification, quantification, analysis) from scratch, employing a new protein database derived from the RNA-Seq analysis performed in this work. This database increased the number and de quality of identifications. A total of 1442 proteins were detected (Table S4), 541 more (Figure S2A), with a better average identification score (Figure S2B) than previous analysis. The higher number of proteins increased the biological meaning of this dataset.

At protein level, the main upregulated pathways (T1 and T3) were phytohormone action, external stimuli response or redox homeostasis (Figure 3) as previously reported (Escandón et al., 2017). Furthermore, stress also increased chromatin reorganization at short times of stress, which may be related to the importance of nuclear and nucleoid processes during stress acclimation revealed at transcriptome level (Figure 2, 3) and/or with the activation of epigenetic mechanisms leading to the acquisition of long-term stress memory as recently described (Lamelas et al., 2020). On the other hand, downregulated pathways are related to cell wall organization, carbohydrate metabolism and the overall secondary metabolism pathway. Metabolome analysis revealed that many secondary metabolites were down-accumulated after stress, but also some antioxidant-related pathways increased (as flavone and flavonol or diterpenoid). Enrichment analyses using GO terms (Figure 3) supported these observations, pointing to hydrogen peroxide or anthocyanin biosynthesis, and other terms coincident with the enrichment at RNA level (Figure 2). Responses at the proteome were delayed from those observed at transcript level, i.e., responses to stress peaked at T3 in proteins and at T1 in transcripts. Protein analyses also pointed to new stress points at endoplasmic reticulum, and the role of RNA methylation and chromatin structure (ASTRA and chloroplast nucleoid) at T1.

**Figure 3:**
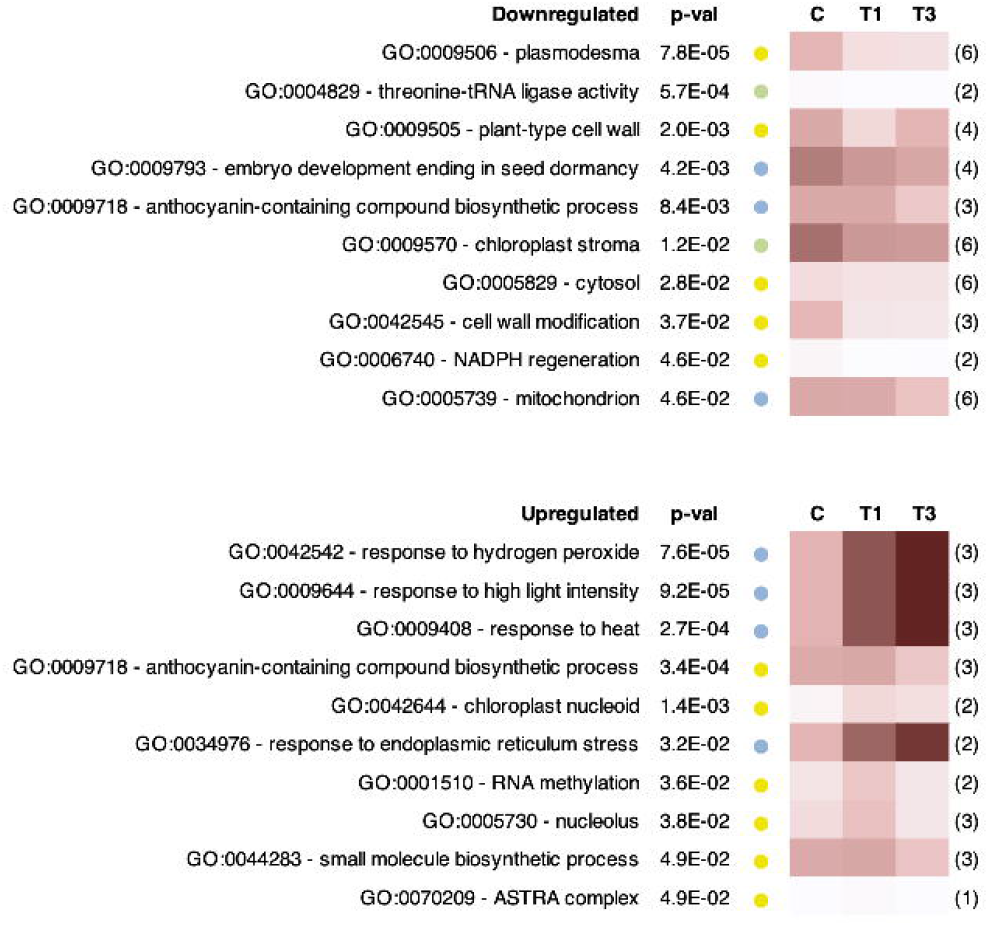
GeneOntology (GO) enrichment analysis of differentially accumulated proteins in response to heat stress at the two sampling times (T1, T3) with respect to controls (C). p-val column indicates the p-values obtained for each term after g:Profiler analyses. Colored dot indicates if term was enriched at T1 (yellow), T3 (blue) or both (green) sampling times. The color of the boxes represents the average expression rate of all differentially expressed genes (p < 0.05; >2-fold change) within each category at each sampling time. The number of differentially expressed genes within each category (TnQ value) is in brackets.

### Different cluster of elements related to time-course stress response were identified by multi-omic data integration

The integration of the transcriptome proteome datasets with available metabolome and physiology datasets (Tables S5, S6; Escandón et al., 2017) allowed a comprehensive overview of heat stress acclimation process in *P. radiata*. A multiblock sPLS-DA (Chen et al., 2021; Singh et al., 2019) was employed for reducing the dimensionality of the datasets and keep only a minimum number of representative variables, and for integrating the different datasets. After this analysis the most important 120 transcripts, 100 proteins, 60 metabolites and 10 physiological parameters were selected (Supplementary Table S7). With these variables the model effectively classified the different samples at each individual omics levels and its combination (Figure 4A). Selected transcripts codify for heat-shock, lipid metabolism (biosynthesis, lipoxygenases), redox homeostasis (GSTU, cytochromes), energy related (various pathways), invertases (cell wall and vacuole), signaling, proteome remodeling (ribosomal units, ubiquitins) and chromatin structure and gene regulation related proteins, including helicases, transcription factors and pentatricopeptide proteins. The presence of chloroplast and mitochondrial pentatricopeptide proteins, reinforced the hypothesis raised after individual analyses that RNA processing in these organelles was a key mechanism for acclimating to stress (Lamelas et al., 2022; Sun and Guo, 2016). The most important proteins for distinguishing the different treatments were different than the transcripts. Heat shock, redox-related proteins (redoxins, S-related proteins), signaling (several 14-3-3, RAS), proteome remodeling related (translation initiation factors), and chromatin structure (histones, helicases) and transcription factors, including NAC proteins. This is not striking as proteome analyses only revealed the most abundant proteins, compared to an almost complete transcriptome analysis. However, despite not keeping transcript-protein pairs, the selected biological functions were conserved. At metabolome level, most of the identified compounds were polyphenols, reinforcing the idea of increased antioxidant mechanisms and hormones. At physiological level, membrane damage (MDA, EL) and hormones (GAs, ABA, iPA) pointing the importance of hormonal signaling, which was not directly revealed by functional enrichment or the presence of receptor or intermediate signalers within transcripts and proteins. Some of the differential transcripts pointed to ethylene and nitric oxide pathways mediating as a part of a complex heat-stress acclimation response, behaving in the same way than other plant species, however these two regulators were not analyzed due to its volatile nature. ABA, overaccumulated in T3, may be acting as thermo-primer mediating the accumulation of ROS scavenging mechanisms, mobilizing sucrose-upregulating invertases, and inducing HSPs (Li et al., 2021).

**Figure 4:**
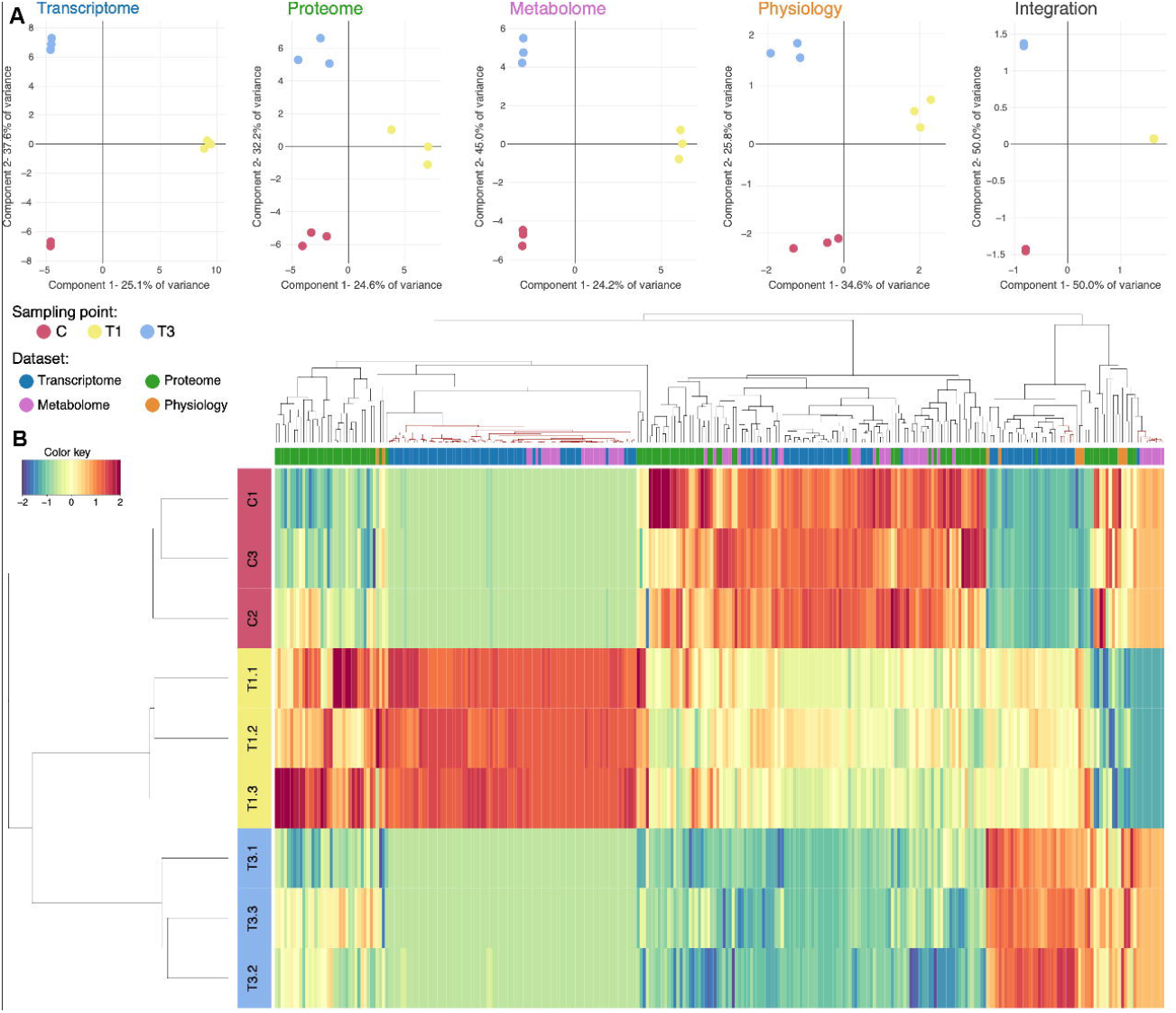
**A**. Sample classification after block analysis of each available dataset and its integration. Bar colors indicate the stress treatment in which this transcript is more expressed (red, control; green, T1; yellow, T3). **B**. Heatmap representation of the most significant variables of the different datasets determined after block-spls analysis. Colors in vertical bar represent the different treatments, while colors in horizontal bar indicate the origin (dataset) of each variable.

The heatmap representation of these variables (Figure 4B) correctly classified the different treatments and showed four main coexpression patterns (overexpressed in C, T1, T1 and T3, and C and T3). In. this heatmap only two clear coexpression cluster can be distinguished at T1 and in C and T3. Interestingly there are several subclusters in T1 cluster, involving light, heat, and redox stress response (PAR1 protein, heat shock protein), gene expression and RNA processing (pentatricopeptide, DNA interacting proteins), and signaling (VAC1). Cluster corresponding of T3 includes several metabolites and hormones. Despite the distance between variables is higher than in previous cases, there is a cluster overexpressed in T1 and T3 which integrates RNA complexes, pentatrichopeptide proteins and a ser/thr kinase which may be regulating retrograde signaling from organelles to nuclei (Lamelas et al., 2022; Sun and Guo, 2016).

### Integrative functional networks revealed the importance of processes related to epigenetic control, protein biosynthesis, photosynthesis, and secondary metabolism

After multi block sPLS-DA analysis two interaction networks were constructed based on distance matrices (Figure 5), or the overlap between distances and biological interactions gathered in STRING/STITCH database (Kuhn et al., 2008; Szklarczyk et al., 2019) (Figure 6) to explain causal relations between the different omic levels. Despite the general preference of networks considering biological information to correct possible mathematical biases (Fukushima et al., 2014), in our hands both kinds of analyses are useful due to the poor functional annotation of Pinus samples. These networks combined the different omic levels and were plot for explaining physiological and metabolomic responses based on the changes at proteome and transcriptome, allowing a comprehensive analysis of all the datasets.

**Figure 5:**
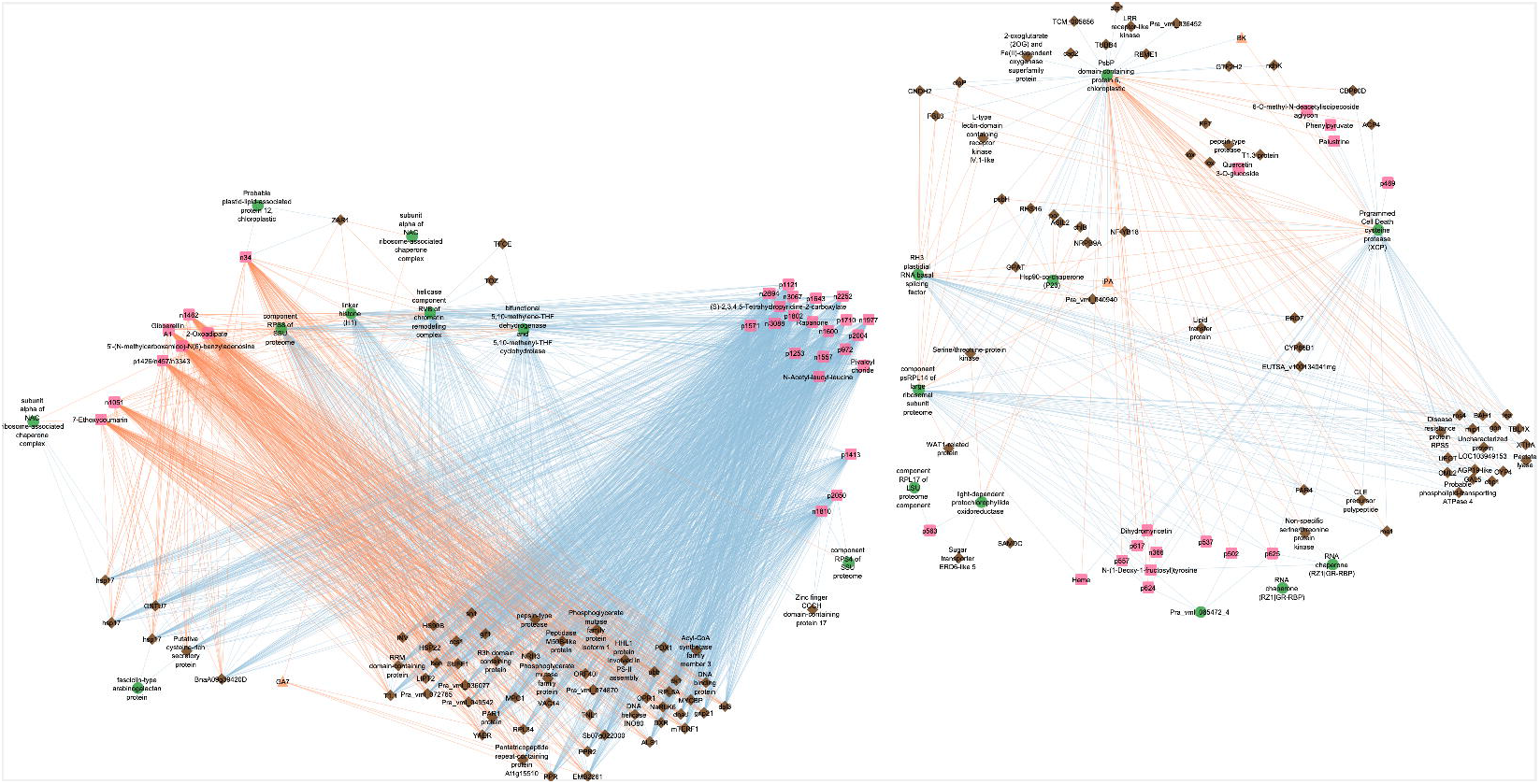
Multiblock-sPLS network constructed with the most significant variables of the different datasets using two thresholds (0.93 to protein connections and 0.99 for the other connections). Physiological parameters are drawn in orange, metabolites in pink, proteins in green and transcripts in yellow. Circles on the network show elements with the same connections.

**Figure 6:**
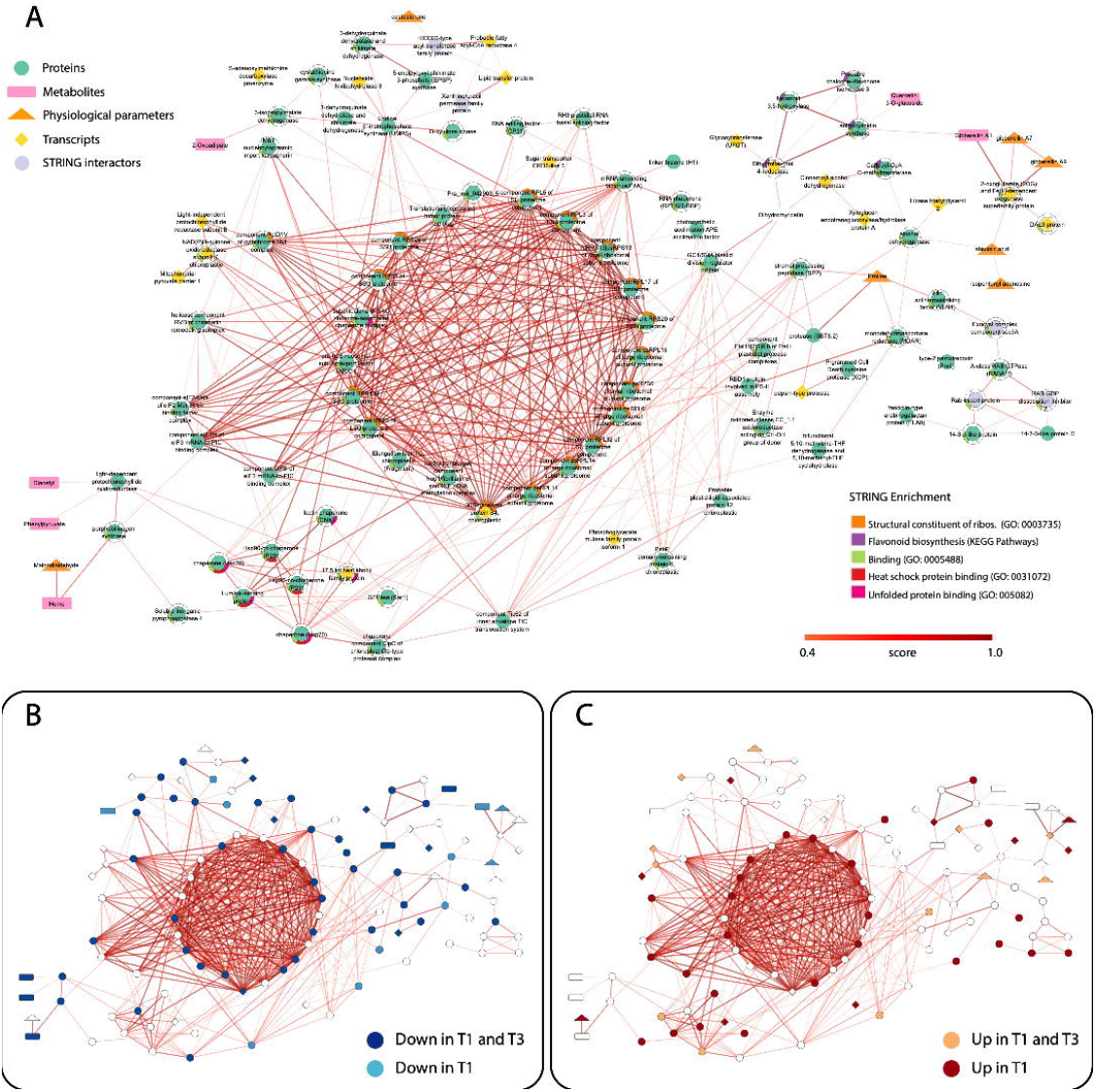
**A**. Resulting STITCH network with the most significant variables of the different datasets determined in block-spls analysis. Physiological parameters are drawn in orange, metabolites in pink, proteins in green and transcripts in yellow. STITCH network was performed using STRING tool and *Arabidopsis thaliana* database. Only interactions scores are equal or higher than 0.4 were represented and for String Enrichment only pathway with a p value < 0.05. In **B** and **C** networks are showing elements down and up regulated across the stress.

As only most important variables were kept at each omic level, the inter and intra omic level interaction networks were very dense, so networks were pruned for showing only most relevant interactions (Figure 6).

### Pentatricopeptide proteins at chloroplast and mitochondria are related to stress signaling and organelle and cell-wide stress responses

Chloroplast pentatricopeptide proteins (Pra_vml_001619, Pra_vml_035514) showed similar interactions, pointing as central players in thermotolerance, being positively correlated with membrane damage (MDA) and with proteins located in chloroplasts such as luminal proteins, prokaryotic ribosomal subunits or chalcone-flavone isomerases. This indicates the role of RNA splicing in stress adaption, but also the retrograde signaling emitted by this organelle (Lamelas et al., 2022), as protein/genes widely expressed across the cell were also correlated as 14-3-3, histone variants, eukaryotic ribosomal subunits, heterogeneous nuclear ribonucleoprotein Q or several NAC proteins (Pra_vml_051671_1, Pra_vml_055001_5) supporting the idea of the cooperative role of the described NAC and pentatricopeptide proteins in stress response processes (Ohbayashi and Sugiyama, 2018), reinforced by its correlation with heterogeneous nuclear ribonucleoprotein Q. Interestingly, chloroplast pentatricopeptide proteins were negatively correlated to sugars (TSS, xylulose kinase, sucrose phosphate phosphatase), gibberelins (GA1, GA7).

On the other hand, mitochondrial pentatricopeptide transcript (Pra_vml_034958) was positively correlated to stress damage (proline content, electrolytic leakage), mitochondrial heat shock proteins, several hormones (RDHZ, BK, iPA and ABA) and a PsP chloroplast protein, while it was negatively correlated to several cell responses and activities. This environmental modulation of mitochondrial responses involving ABA and pentatricopeptide proteins have been already described for Arabidopsis (Liu et al., 2010).

### The inclusion of a functional information layer revealed a complex cellular coordination from organelle transcription to hormone-signaled systemic response

Network including functional interactions (Figure 6), missed some of the correlations shown above due to poor annotations, but showed the central role of proteome remodeling with several ribosomal subunits, regulatory elements (ARX1, NAC, snoRNP-rRNA methylation) and translation factors conforming a central core. This core relates to seven HSP/chaperones through ARX1 and snoRNP rRNA methylation complex. The interaction of these elements points to the nucleolus and nuclear pores as hotspots for adapting to temperature change. In nucleolus, NAC may interact with nuclear pentatricopeptide, causing alternate processing of RNAs (Ohbayashi and Sugiyama, 2018) and the formation of RNA methylation complexes with snoRNP (Wu et al., 2021) which exports stress adapted rRNA forms through ARX1 (Beine-Golovchuk et al., 2018) to the cytoplasm. Interestingly this central core was also correlated to several RNA chaperones, splicing or unfolding factors reinforcing the previous hypothesis about the importance of differential RNA processing and methylation during stress acclimation and supporting the functional enrichments (Figure 2 and 3) obtained after analyzing whole datasets. A long-term monitoring of some of these genes demonstrated that the increased transcriptional activity ends when acclimation was achieved as RH3 plastidial DEAD-BOX helicase, Helicase RVB, RZ1 RNA chaperone, or ribosomal RPS4 reduced its expression to control or below-control levels after five days of stress exposure (Figure S3).

A second interaction center was defined around proline, a common stress marker in trees, was linked to several proteases and peptidases, so its accumulation can be the direct results of protein degradation during proteome remodeling processes. Proline was linked to a cluster of signaling-related proteins comprised of 14-3-3, proteins which accumulates under stress and appear to be induced by ABA and modulate a wide array of targets (Abreu et al., 2013) through a bridge of RAB proteins, and through villin-actin and SEC3A, related to cytoskeleton structure and function. As RAB proteins mediates not only several stress responses (El-Esawi and Alayafi, 2019) but also regulate vesicle trafficking results (Stenmark, 2009), it seems that remodeling intracellular trafficking is also necessary for stress acclimation in pine needles.

Hormones (ABA, iPA, Gas 1,7,9) interaction center was connected to flavonoid metabolism and to alcohol dehydrogenase, pointing the reduction of oxygen within plant cells in response to oxidative stress.

A small cluster of shikimic acid related enzymes, recently related to stress response via programmed cell death (Lu et al., 2022), was linked to UMP synthase and to lipid related transcripts (Lipid transfer protein, Acyl carriers and transferases) suggesting the importance of this pathway during the remodeling of the fatty acids. Shikimate seems to be also related to epigenetics as is linked to S-adenosylmethionine metabolism (NRH3 and SAMDC transcripts were upregulated after stress) through UMP synthase.

## Conclusions

The assembly of a specific transcriptome, and its combination with other omics, allowed an unprecedented understanding of the mechanisms behind acclimation, involving complex interrelated processes from molecular to physiological level. Specific nucleolar or nucleoid activities seems to be the key acclimating process, producing specific RNA isoforms and participating in anterograde-retrograde stress signaling. This mechanism was connected by specific proteins or transcripts to already known heat stress-response mechanisms such as redox, heat shock proteins or ABA mediated, but also to new candidates such as shikimic pathway, photosynthetic pigments, or proline centric proteases activities, effectively linking specific physiological and biochemical responses to specific molecular/regulatory elements. Despite more efforts are needed to fully understand heat stress acclimation in pines, this work provides molecular evidence about the central mechanisms that underlaying response to heat stress in conifers.

## Data Availability Statement

The raw RNA-seq data have been deposited with links to Bioproject accession number PRJNA851373 in the NCBI Bioproject database (https://www.ncbi.nlm.nih.gov/bioproject/). The mass spectrometry proteomics data including RAW, msf and pepXML files have been deposited to the ProteomeXchange Consortium via the PRIDE (Perez-Riverol et al., 2022) partner repository with the data set identifier PXD032754. The metabolomics data have been deposited to the Metabolights repository (https://www.ebi.ac.uk/metabolights/) with the data set identifier MTBLS4567.

## Acknowledgements

Funding for this work was provided by the Spanish Ministry of Economy and Competitiveness (AGL2016-77633-P and PID2019-1071076B-100 projects).

## Supplementary Material

**Supplementary Figure 1**. Experimental design, indicating the different omics layers analyzed at each point: Phy, physiological parameters and hormonal analysis; Met, Metabolome; Prot, proteome; Trans, transcriptome and Target Trans, targeted transcriptomics. C, control; T1/2, 3 h after 40 °C on day 1; T1, 6-h heat exposure on day 1; T2, day 2; T3, day 3; T5, day 5.

**Supplementary Figure 2**. Reanalysis and identification of the protein dataset using the in-house database created with the RNA-seq data. A) Identification score plot comparing the new identifications with those previously obtained in Escandon et al. (2016), B) Venn diagrams and C) Heatmap with protein re-identification using Mapman.

**Supplementary Figure 3**. Selected candidates present in both interaction networks (DIABLO and STITCH) analyzing their tendency at the proteomic and transcriptomic level. Transcriptomic level has been included at a massive level (RNA-seq) as well as directed (RT-qPCR) by extending its trend with the intermediate treatment T2 and further T5.

**Supplemental Tables will be provided in accepted version of the Manuscript**

