## Supplementary figures and images for "Multiomics analyses reveal the central role of nucleolus and nucleoid machinery during heat stress acclimation in *Pinus radiata*"

### Figure S1.png

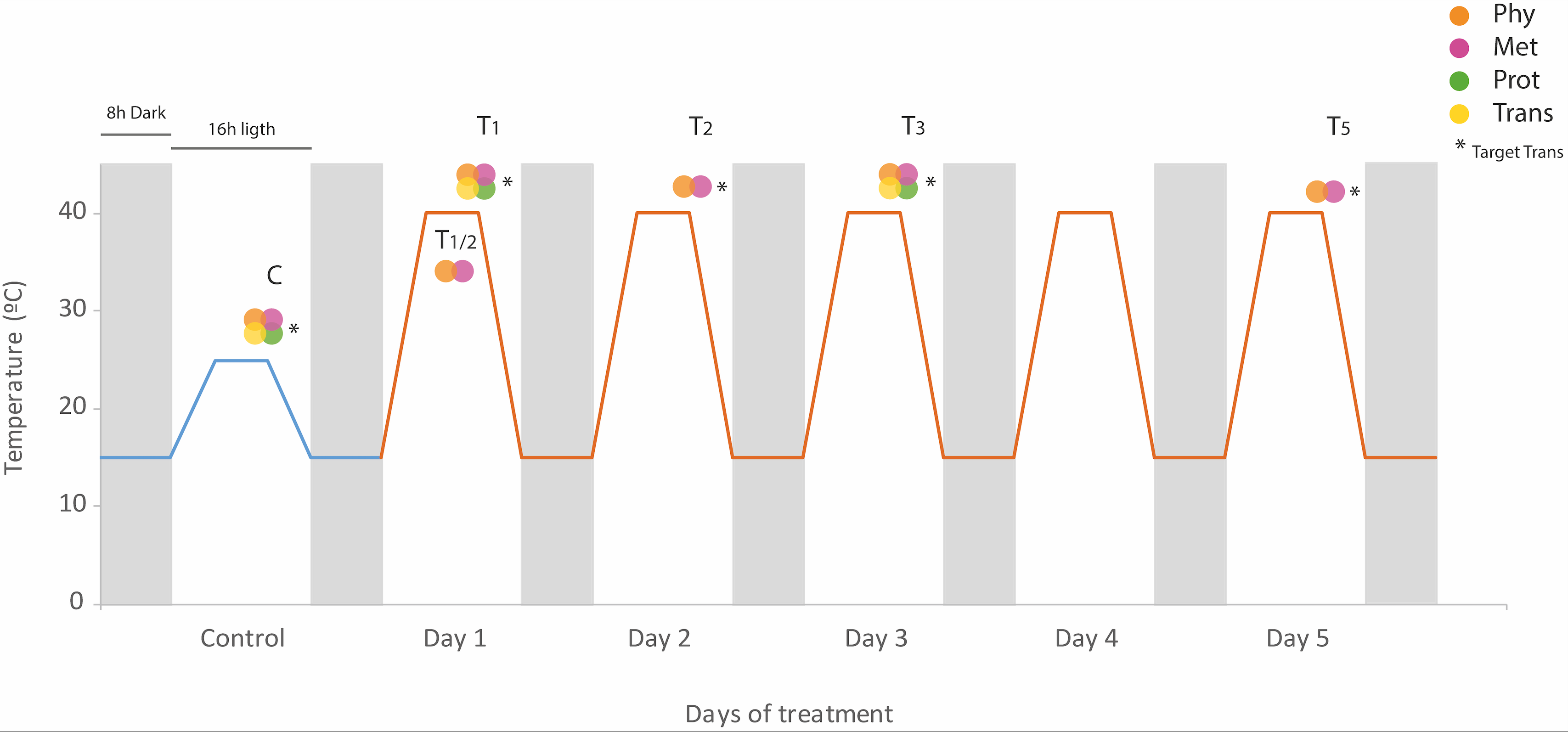

### Figure S2.png

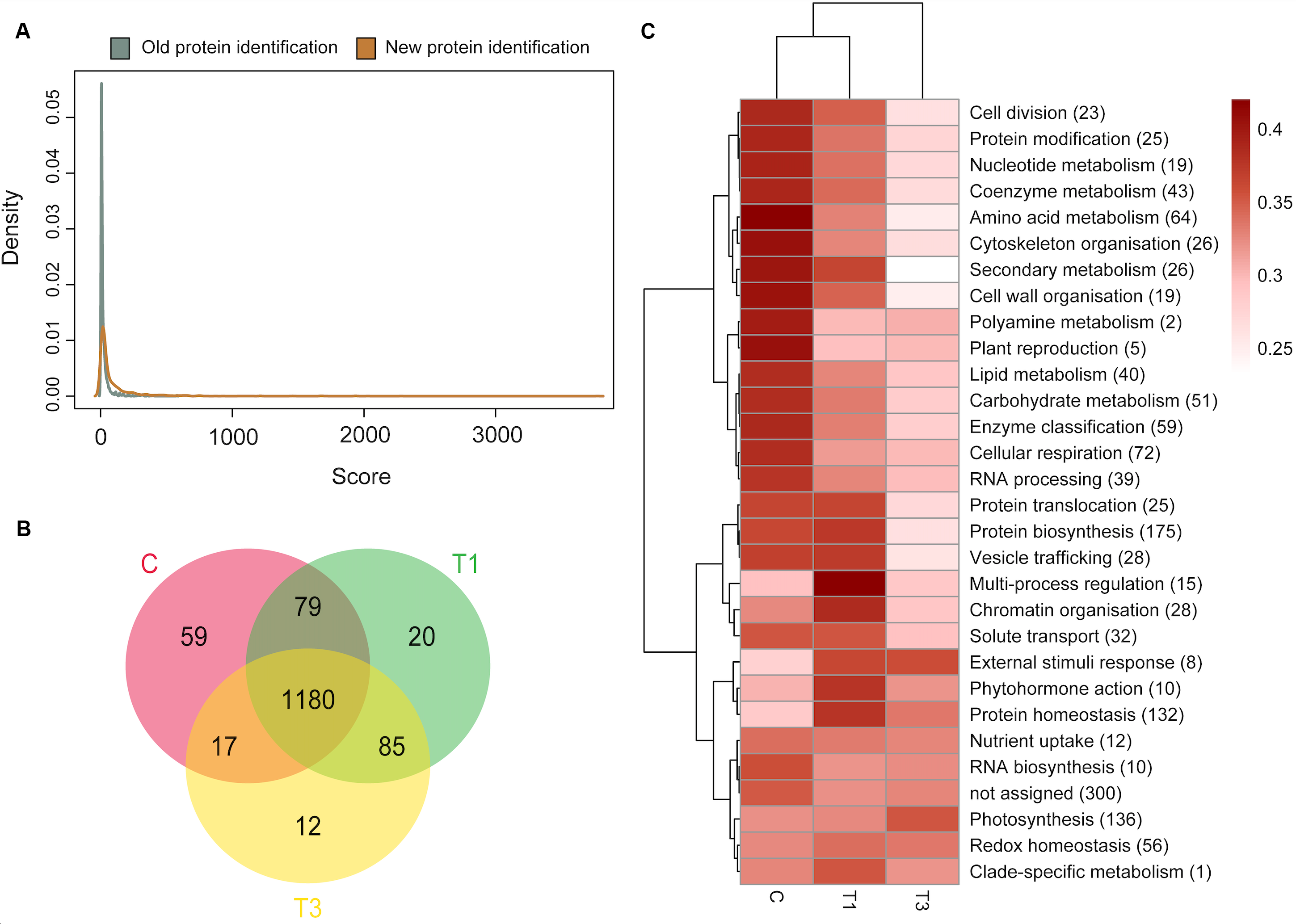

### Figure S3.png

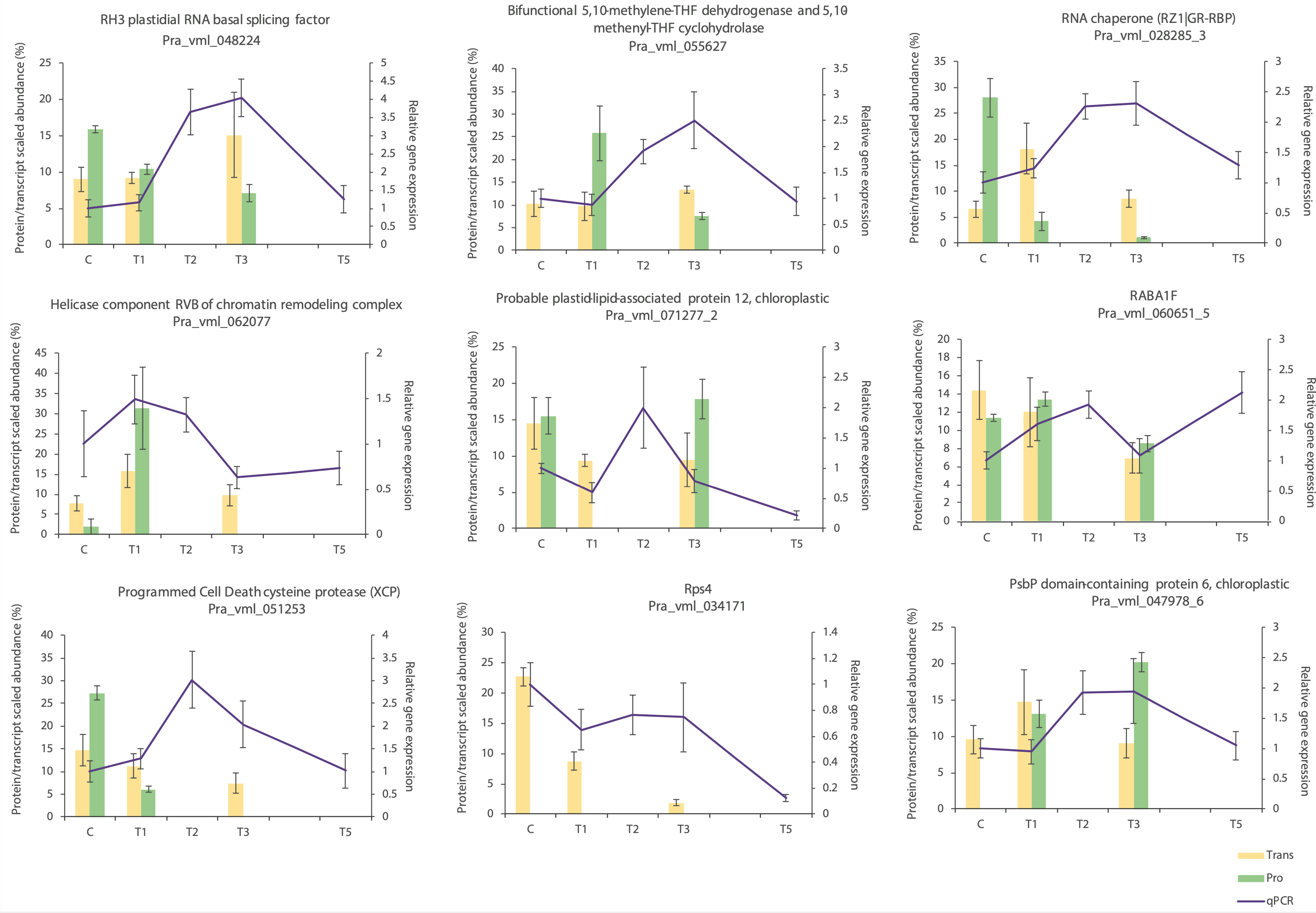
